# Molecular mechanisms of 10-Butyl Ether Minocycline (BEM), a novel non-antibiotic tetracycline, as a potential treatment for inflammatory and neuroimmune-related disorders

**DOI:** 10.1101/2025.06.23.661183

**Authors:** Abdul A. Shaik, Praneetha Panthagani, Xiaobo Liu, Stephany Navarro-Turk, Jeremy Garza, Monica Aguilera, Jordan Sanchez, Kushal Gupta, Abdul Hamood, Ted W. Reid, Bruce Blough, Elliott Pauli, Jeremy D. Bailoo, Susan E. Bergeson

**Affiliations:** Department of Cell Biology and Biochemistry, School of Medicine, Texas Tech University Health Sciences Center, Lubbock, TX, United States; Department of Pharmacology and Neuroscience, School of Medicine, Texas Tech University Health Sciences Center, Lubbock, TX, United States; Department of Pathology, School of Medicine, Texas Tech University Health Sciences Center, Lubbock, TX, United States; Department of Host-Microbiome Interactions, St. Jude Children’s Research Hospital Memphis, TN, United States; Department of Immunology and Molecular Microbiology, School of Medicine, Texas Tech University Health Sciences Center, Lubbock, TX, United States; Department of Ophthalmology & Visual Sciences, School of Medicine, Texas Tech University Health Sciences Center, Lubbock, TX, United States; Discovery Sciences, RTI International, Research Triangle Park, North Carolina, United States; Early to Late Stage L.L.C, North Carolina, USA; McGovern Medical School, The University of Texas Health Sciences Center, Houston, TX, United States

## Abstract

The pleiotropy of minocycline (MINO), including anti-inflammatory, antioxidant, anti-migratory, anti-MMP, and neuroprotective effects, has been extensively reported. A novel non-antibiotic minocycline derivative, 10-butyl ether minocycline (BEM), was synthesized to retain the pleiotropy of minocycline while minimizing side effects such as antibiotic resistance and gut dysbiosis. Previously, we showed that BEM reduced alcohol consumption in dependent murine and porcine models of Alcohol Use Disorder (AUD). In this study, we investigated the molecular mechanisms of BEM to determine its potential as a therapeutic agent for neuroimmune and inflammatory conditions such as AUD. Here, we report that BEM showed a nearly complete loss of antimicrobial activity against *E. coli*, *S. typhi,* and *C. albicans*. BEM showed a dose-dependent reduction in cell viability as measured by the MTT assay, similarly to MINO. BEM also suppressed LPS-induced microglial activation as shown by reduced Iba1 expression in immunohistochemistry and western blot analyses. Inhibition of MMP-9 by BEM (IC50 = 42.2 µM) was improved compared to MINO (IC50 = 60.3 µM) while MMP-8 inhibition was moderate (IC50: BEM = 69.4 µM; MINO = 45.4 µM). BEM was found to be effective in inhibiting VEGF-induced endothelial cell migration and L-glutamine-induced ROS levels. Limited inhibition of 15-LOX activity was observed (IC50: BEM = 92.6 µM; MINO = 65.6 µM). BEM was not toxic to mitochondria, even at high concentrations (200 µM). By eliminating antimicrobial properties while preserving therapeutic pleiotropy, BEM presents an advancement in the development of a promising candidate with multimodal mechanisms to treat neuroimmune-inflammatory pathologies.

**Impact:** Our current approach to correlate non-antibacterial actions of BEM to MINOs established mechanistic effects will enable the informed use of BEM for several medical indications including for inflammation and neuroimmune conditions. The focus on BEM’s multimodal actions and long-term safety during drug discovery represents a paradigm shift toward complex therapeutic drug development and repositioning, improving upon traditional singular high-affinity target-based approaches. Such new drug discovery attempts could potentially enhance treatment relevance in complex disorders with multiple targets and theoretically guide the creation of second-generation analogs.

**Significance statement:** We report mechanisms of action for BEM, a minocycline analog under evaluation for the treatment of Alcohol Use Disorder, which may also show efficacy for other complex disease processes that involve inflammatory or neuroimmune components. We show that BEM had a nearly complete loss of antimicrobial action, yet retained the pleiotropy of MINO, likely making it a better multimodal therapeutic for long-term treatment of complex diseases with neuroimmune-related components.

**Visual abstract:** 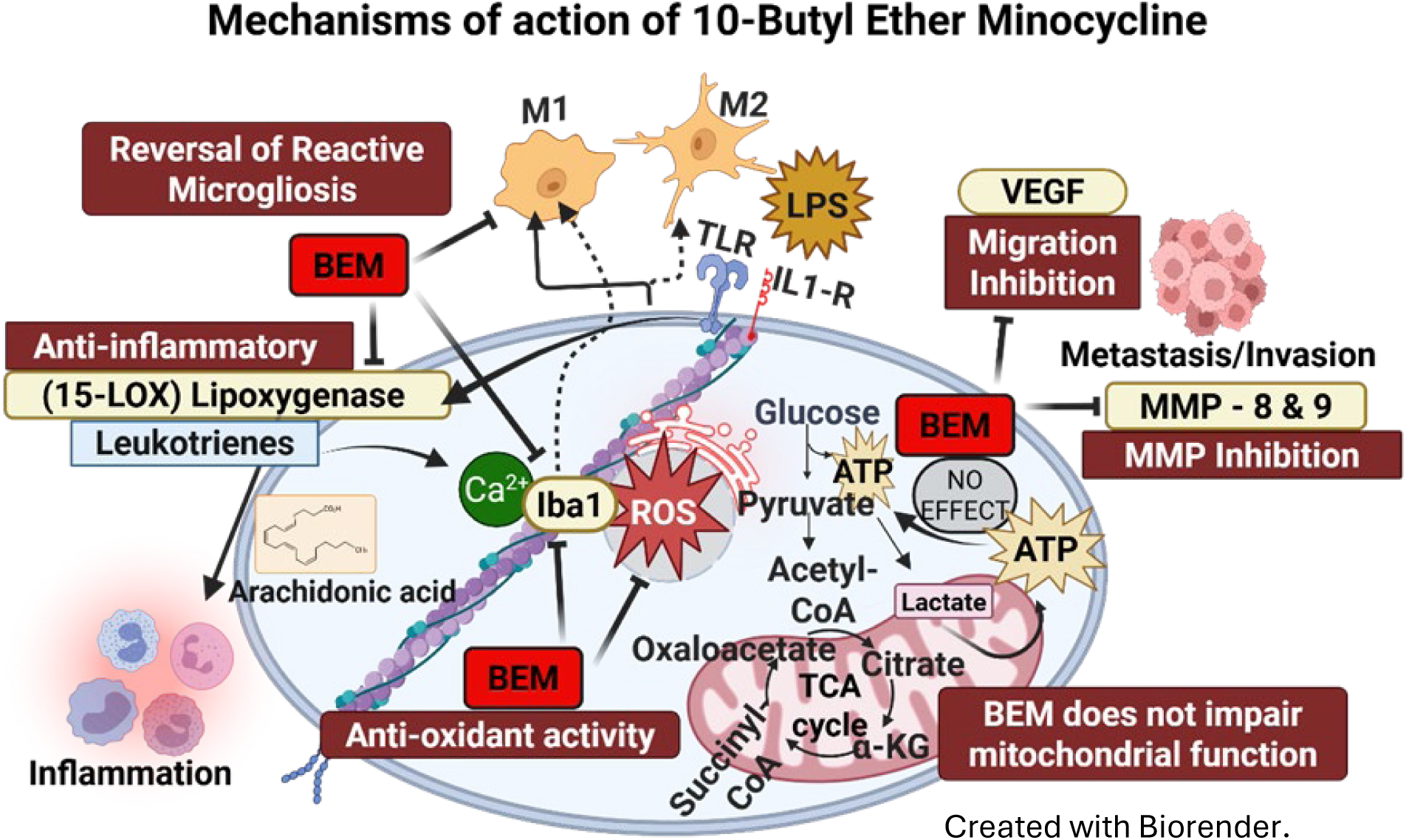

## Introduction

### Background

Tetracyclines are members of a group of broad-spectrum antibiotics which are effective against gram–positive and –negative bacteria and protozoan parasites (Chopra & Roberts, 2001). They have been used in clinical settings for over 85 years for the treatment of acne (Meynadier & Alirezai, 1998), Lyme disease (Nowakowski et al., 1995), and urinary related infections (Anderson et al., 1966), to name a few. The primary side effect from the use of tetracyclines is gut dysbiosis, which can take the form of loss of beneficial bacteria, overgrowth of harmful bacteria, and loss of diversity in the gut flora (Patangia et al., 2022). The primary target of tetracyclines is the 30S (Brodersen et al., 2000). However, the mammalian mitochondrial ribosome is generally agreed to be evolutionarily related (Geiger et al., 2023) and tetracycline binding leads to inhibition of mitochondrial protein translation (Moullan et al., 2015; Mortison et al., 2018).

Tetracyclines interact with diverse non-antibiotic cellular targets, receptors and signaling pathways, indicating a pleiotropic pharmacological profile. Contemporary research highlights their potential to modulate neuroimmune pathways via off-target mechanisms, expanding their therapeutic scope beyond infectious diseases (Asadi et al., 2020; Park et al., 2020; Lopez-Navarro & Gutierrez, 2022; Markulin et al., 2022; Rahmani et al., 2022; Sanshita et al., 2023; Kounatidis et al., 2024; Ritter et al., 2024). This growing body of evidence has motivated extensive repurposing efforts of tetracyclines for central nervous system and immune-inflammatory indications where pathologies are driven by multi-modal mechanisms (Elewa et al., 2006; Garrido-Mesa et al., 2013; Chaves Filho et al., 2021; Garrido-Mesa et al., 2022; Markulin et al., 2022).

### Repurposing Minocycline

Minocycline (MINO), a common, second generation, semi-synthetic tetracycline antibiotic, has been widely studied in drug repurposing efforts for complex multi-factorial disorders due to its neuroimmune modulatory, anti-migratory, and anti-inflammatory off-target effects (Homsi et al., 2010; Kamal et al., 2020; Moura et al., 2022). Several mechanisms underly MINO’s off-target effects and include: 1) its neuroprotective effects are associated with a reduction in microglial activation (Wang et al., 2018), and it exerts 2) anti-inflammatory actions in part through inhibition of microglial proliferation (Yrjänheikki et al., 1998; Yew et al., 2019; Du et al., 2022), 3) attenuation of lipopolysaccharide (LPS)–induced neuroinflammation (Henry et al., 2008; Qaid et al., 2022), and 4) suppression of arachidonate 5-lipoxygenase (5-LOX) expression and enzyme activation after focal cerebral ischemia (Chu et al., 2007). The anti-proliferative action of MINO could potentially be mediated via inhibition of vascular endothelial growth factor (VEGF) (J. S. Yao et al., 2004; Yao et al., 2006; Higashi et al., 2018). MINO had also demonstrated strong matrix metalloproteinase (MMP) inhibitory effects (Switzer et al., 2011; Chang et al., 2017; Vandooren et al., 2017; Vellimana et al., 2017) that are known to affect VEGF-induced cell migration (Ramamurthy et al., 2002; Yao et al., 2004; Machado et al., 2006; Lee et al., 2007; Chang et al., 2017). In addition, MINO suppressed oxygen radical formation due to direct antioxidant effects are comparable to alpha-tocopherol, independent of Fe^2+^ chelation (Jakub et al., 2021; Rok et al., 2021). This antioxidant activity of MINO had been attributed to its multi-substituted phenol ring structure, which resembles vitamin E (Kraus et al., 2005a).

MINO has been evaluated as a repurposed therapeutic for the treatment of several disorders and disease states. In anxiety and depression, stress-induced neuroinflammation is implicated in disease progression, and MINO has shown promise in mitigating these effects (Liu et al., 2018; Zhang et al., 2019; Camargos et al., 2020; Gajbhiye et al., 2022; Iglesias et al., 2024). MINO has been found to be beneficial as an add-on therapy for treatment resistant major depressive disorder and bipolar depression (Murrough et al., 2018; Nettis et al., 2021). MINO has been shown to alleviate methamphetamine (MA) withdrawal symptoms (Fujita et al., 2012), attenuate the maintenance and reinstatement of drug-seeking behavior (Attarzadeh-Yazdi et al., 2014; Ghavimi et al., 2022) and has shown clinical promise in a reported human case study of MA–use disorder (Tanibuchi et al., 2010). In a recent pilot study, MINO was found effective as an adjunctive treatment for MA-induced psychosis and neuropsychological impairments in five patients with treatment resistant MA use disorder (Alavi et al., 2021). Altogether, these findings highlight MINOs potential use as a repurposed therapeutic candidate for conditions involving neuroimmune dysregulation and neuroinflammation.

### Treatment of Alcohol Use Disorder (AUD) with Minocycline

AUD is a complex trait with numerous genetic and environmental influences including neuroimmune dysregulation and neuroinflammation. Our lab and others have found that long-term alcohol consumption induces microglial-related transcriptome changes, associating groups of genes involved in immune regulation and voluntary alcohol consumption (Mulligan et al., 2006a; Mulligan et al., 2008; Blednov et al., 2011; McCarthy et al., 2017). Through bioinformatic pathway analyses we identified neuroimmune signaling as a key mechanism underlying AUD pathology (Mulligan et al., 2006b; Agrawal et al., 2011; Agrawal et al., 2014). Since then, therapeutic targeting of neuroimmune signaling in AUD is now widely recognized as an exciting new research avenue (c.f., Erickson et al., 2019 for review).

Building on the expanding therapeutic potential of tetracyclines, our lab was the first to identify MINO as a potential therapeutic candidate for AUD (Agrawal et al., 2011) due to its well-documented anti-inflammatory (Dunston et al., 2010; Schmidt et al., 2021), neuroprotective (Elewa et al., 2006; Plane et al., 2010; Ortiz et al., 2020), and immune-modulatory properties (Dutta et al., 2010; Garrido-Mesa et al., 2011; Schmidt et al., 2021). MINO crosses the blood brain barrier and reduces microglial activation, a driver of neuroinflammation (Gajbhiye et al., 2018). In mice, MINO reduced voluntary alcohol intake, mitigated withdrawal-related anxiety, and decreased alcohol cue-induced relapse. In male Wistar rats, MMP-9 inhibition in central amygdala prevented acute withdrawal-induced escalation of alcohol self-administration (Go et al., 2019). MMPs drive plasticity dependent learning by remodeling the extracellular matrix. Escalation of alcohol self-administration and addiction is MMP-dependent (Yin et al., 2020). In AUD pathophysiology, neuroimmune interactions, specifically in microglia are known to play an important role (Li et al., 2024; Rasool et al., 2024). Chronic inflammation in AUD can lead to compromised blood brain barrier integrity, invasion of peripheral immune cells, mitochondrial dysfunction and dampened neurogenesis, all of target features which may be targeted by MINO (Portis & Haass-Koffler, 2020).

### Development of Minocycline analogs

We proposed that the off-target pharmacology of MINO holds potential for treating inflammation and neuroimmune conditions, such as AUD. Our previous studies have demonstrated the preclinical efficacy of semi-synthetic tetracyclines, including MINO, in reducing alcohol consumption in rodent models (Agrawal et al., 2011; Martinez et al., 2016; Syapin et al., 2016). However, as is the case with all tetracyclines, its prolonged use was limited by the risk of gut dysbiosis (Matthew et al., 2023), and, to some extent, the potential inhibition of mitochondrial translation, raising concerns about long-term clinical safety and compliance.

To address this limitation, based, in part, on our tetracycline screen for efficacy to reduce alcohol consumption, which elucidated Structure Activity Relationships (Syapin et al., 2016), and the solved structure of tetracycline bound to the bacterial ribosome (Schedlbauer et al., 2015) we developed a series of MINO synthetic analogs, aiming to retain its off-target therapeutic benefits and increase CNS penetration while reducing antibiotic-associated side effects. 10-Butyl Ether Minocycline (BEM), our lead molecule, is both chemically and pharmacologically novel. Chemically, the C-10 butoxy group in 10- butyl ether minocycline (BEM) introduces steric bulk at a ribosome-contacting position, decoupling antibacterial binding while retaining the divalent ion chelating/phenolic features (e.g., C11/C12 β-diketone, C7/C4 dimethyl amino) thought to underlie MINOs pleiotropy. Pharmacologically, BEM shows favorable non-antimicrobial, physicochemical– and pharmacokinetic– toxicological profiles, most of which we report here for the first time. Overall, we hypothesized that the off-target mechanisms contributing to MINOs efficacy in treating high alcohol consumption, may contribute to BEMs effectiveness in treating AUD.

We successfully demonstrated BEM’s efficacy in reducing alcohol consumption in alcohol dependent C57BL/6J mice subjected to chronic intermittent ethanol exposure (Panthagani et al., 2024). In a minipig model of alcohol dependence, where animals were diagnosed with severe AUD based on DSM-V-TR criteria, we again showed that BEM significantly reduced alcohol consumption (Liu et al., 2023; Liu, Gutierrez, et al., 2024; Liu, Panthagani, et al., 2024). These results therefore established BEM’s therapeutic efficacy in both mice and porcine AUD models. We have also showed that other MINO derivatives like diacetyl minocycline, can mitigate pathological choroidal neovascularization in a mouse model, potentially through matrix metalloproteinase (MMP-9) inhibition (J. Willms et al., 2023).

In the current study, we explore the mechanisms likely underlying BEM’s efficacy in treating AUD and potentially other complex diseases and disorders which share similar underlying etiology. We hypothesized that BEM had retained the pharmacodynamic pleiotropy of MINO while losing its anti-microbial activity. We drew on existing MINO-related literature and experimental findings of BEM to establish their shared, and distinct, underlying mechanisms of action.

## Materials and methods

### 10-Butyl Ether Minocycline

Our lead molecule, BEM·HCl·3H₂O, is a MINO analog with the IUPAC nomenclature (4S,4aR,5aS,12aR,E)-2-(amino(hydroxy)methylene)-10-butoxy-4,7-bis(dimethylamino)-11,12a- dihydroxy-4a,5a,6,12a-tetrahydrotetracene-1,3,12(2H,4H,5H)-trione. Pharmaceutical grade BEM was synthesized as a hydrochloric acid salt by Curia Global Inc. (Italy) under good manufacturing practices conditions, at a purity of ≥ 97 % (synthesis scheme undisclosed). BEM has a chemical formula of C₂₇H₃₇Cl₂N₃O₇·3H₂O and a molecular weight of 622.55 g/mol. BEM’s stability testing at Curia Global Inc. under conditions of 25 °C / 60 % RH (relative humidity) and 40 °C / 75 % RH for 15, 30, and 45 days, did not show significant changes in purity. A re-test conducted after 1, 3, 6 and 10 months of storage at -20 °C again revealed no impurities. As stated in the certificate of analysis from Curia Global Inc., BEM is a yellow solid that meets identification criteria via infrared spectroscopy, nuclear magnetic resonance and high-performance liquid chromatography–mass spectrometry. The chemical structures of MINO and BEM are shown in Figure 1.

**Figure 1.**
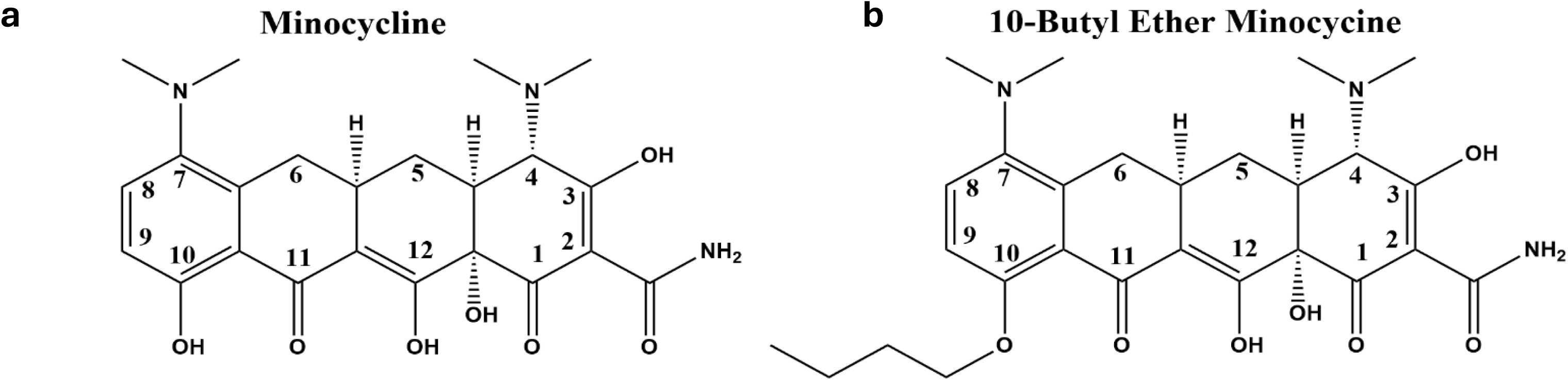
The structure of (a) minocycline (MINO) and its analog (b) 10-butyl ether minocycline (BEM). A butoxy substitution at the C10 position was introduced to reduce or eliminate BEM’s antibiotic activity.

### Antibacterial Assay (Filter Paper Disc Method)

The antibacterial activity of BEM and MINO against *Escherichia coli* (*E. coli*) and *Salmonella typhi* (*S. typhi*) was evaluated using the filter paper disc method (Balouiri et al., 2015). Both *E. coli & S. typhi* were streaked onto individual lysogeny broth (LB) agar plates (Fisher Bioreagents, Mexico) to ensure uniform bacterial growth (Sanders, 2012). Immediately after streaking, cellulose discs infused with varying concentrations of BEM and MINO (1.25, 3.125, 6.25, 12.5, 25, and 50 µg/mL) were placed in designated quadrants on an agar plate. The plates were then incubated overnight at 37 °C, and antibacterial activity was assessed by measuring the zone of inhibition (ZOI), i.e., the diameter of the region with no bacterial growth around the filter paper disc.

### Antifungal Activity

The minimum fungicidal concentration of BEM was evaluated following established protocols (Bertout et al., 2011; Alastruey-Izquierdo et al., 2015; J. O. Willms et al., 2023). Briefly, BEM and MINO were diluted to final concentrations of 0, 25, 50, and 100 µg/mL in yeast peptone dextrose (YPD) broth (Fisher Bioreagents, Mexico) and inoculated with *Candida. albicans* (*C. albicans)* (ATCC 3147, Pamela Parr) at a density of 10⁵ colony-forming units (CFU)/mL. The cultures were incubated at 35 °C for 24 hours, after which they were serially diluted, 10-fold, and plated onto YPD agar. CFU’s were manually counted to assess fungal viability.

### Cell Viability Assay

The safety profile of BEM was assessed using a neuroblast (NB) cell viability assay and compared to the parent compound, MINO, utilizing the 3-(4,5-Dimethylthiazol-2-yl)-2,5-Diphenyltetrazolium Bromide (MTT) (Thermo Fisher, Germany) assay. Briefly, NB cells were seeded in a 96-well plate and cultured to 70% confluency in Dulbecco’s Modified Eagle Medium (DMEM) (Gibco, St. Louis, MO) supplemented with fetal bovine serum (Gibco, St. Louis, MO). Cells were incubated overnight at 37 °C with 5% CO₂. Once confluent, BEM and MINO were administered at concentrations of 0.1, 1, 10, 50, 125, 250, and 500 µM, and cells were incubated for 24 hours at 37°C. Following treatment, the drug-containing DMEM media was replaced with phenol red-free media, and MTT solution was added to each well to reach a final concentration of 0.5 mg/mL. After a 2-hour incubation, 100 µL of DMSO was added to dissolve the formazan crystals, and the plate was placed on a shaker to complete solubilization. Absorbance, measured at 590 nm within one hour, was used to determine cell viability. The resulting absorbances were used to calculate IC50 using a log(inhibitor) vs. response (three-parameter) model in GraphPad Prism 10.5.0.

### Matrix Metallo Proteinases (MMP-8/9) Inhibition Screening Assay

The inhibitory activity of BEM and MINO against MMP-8 and MMP-9 was assessed by measuring the percentage inhibition at various concentrations. For MMP-9, the concentrations tested were 20, 40, 60, and 80 µM, while for MMP-8, the concentrations were 30, 60, 90, and 120 µM. The results were compared to the prototypic inhibitor N-Isobutyl-N-(4-methoxyphenylsulfonyl)Glycyl Hydroxamic Acid (NNGH). The colorimetric assay was conducted in a 96-well microplate format using a chromogenic substrate (Ac-PLG-[2-mercapto-4-methyl-pentanoyl]-LG-OC2H5). According to the MMP-8/9 inhibitor screening assay colorimetric kit (Abcam, MMP-8 - ab139452, MMP-9 - ab139448, Waltham, MA), BEM, MINO, and the reference inhibitor NNHG were added to the assay buffer followed by the MMP enzymes. The reaction mixture was incubated at 37 °C with 5% CO₂ for 60 minutes. After incubation, the chromogenic substrate was added, and absorbance (ΔA) was measured every minute, for 10 minutes, at 412 nm. The percentage inhibition of MMP-8/9 for each compound was calculated as follows:

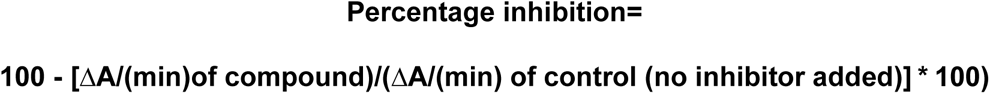

The resulting dose-dependent percentage inhibition values of MINO and BEM were used to perform nonlinear regression analysis and fit to a log(inhibitor) vs. response – variable slope (four parameters) model. The slopes of BEM and MINO were then compared using simple linear regression in GraphPad Prism 10.5.0.

### Microglial Activation Assay

Immortalized murine N9 microglial cells were cultured using the Nunc Lab-Tek II chamber slide system (Thermo Fisher Scientific, Waltham, MA) and incubated overnight at 37 °C with 5% CO_2_ in DMEM media to reach 70% confluency. The cells were then treated with 0.1 µg/mL lipopolysaccharide (LPS; Invitrogen, Cat No 00497693) for 12 hours, followed by treatment with BEM and MINO at a 20 µM concentration for an additional 12 hours. After treatment, the cells were washed with phosphate buffer saline and fixed in 4% paraformaldehyde for 10 minutes. The fixed cells were permeabilized with a solution of 95% ethanol and 5% glacial acetic acid for 5–10 minutes.

The cells were incubated overnight with a 1:100 dilution of Iba1-specific goat polyclonal antibody (Abcam, ab178847), followed by a 4-hour incubation with a 1:100 dilution of anti-goat secondary antibody conjugated with fluorescein (FITC; Santa Cruz Biotech, sc-2356). The nuclei were stained with ProLong™ Gold Antifade Mountant with DAPI (Thermo Fisher Scientific, Waltham, MA). Images were captured using a Zeiss Axio Imager M2 confocal microscope (60X, oil immersion). Immunohistochemistry was performed to assess microglial activation. The change in morphology of the LPS treated cells, compared to control, formed the basis of degree of ramification for all the treatments. The %ramified cells were manually calculated in each treatment and graphed using GraphPad Prism 10.5.0.

To further validate the immunohistochemical findings, N9 microglial cells were cultured in 6-well plates and allowed to grow overnight at 37 °C with 5% CO_2_ to 70% confluency in DMEM media. The cells were treated with 25 ng/mL LPS for 12 hours, followed by 25 µM BEM and MINO for an additional 12 hours. The treatment was terminated by removing the drug-containing media, and the cells were washed three times with PBS. Protein extraction was performed by lysing the cells with RIPA lysis buffer (Thermo Fisher Scientific, Waltham, MA) containing Halt Protease Inhibitor Cocktail 1X™ (Thermo Fisher Scientific, Waltham, MA). The lysates were collected and centrifuged, and the supernatants were analyzed using the Pierce™ BCA Protein Assay Kit (Thermo Fisher Scientific, Waltham, MA). Protein separation was done via electrophoresis using a 12% Criterion™ TGX Precast Midi Protein Gel (Bio-Rad Laboratories, Hercules, CA).

The proteins were transferred onto an Immun-Blot® PVDF Membrane (Bio-Rad Laboratories, Hercules, CA), which was then probed overnight with primary antibodies: Anti-Iba1 (Abcam, ab5076) and anti-β-Actin rabbit monoclonal antibodies (Cell Signaling, β-Actin (13E5) Rabbit mAb #4970) (1:100 dilution). The next day, anti-rabbit HRP-conjugated secondary antibody (Cell Signaling, Anti-rabbit IgG, HRP-linked Antibody #7074) (1:100 dilution) was applied, and the blot was developed using SuperSignal West Pico PLUS Chemiluminescent Substrate ECL (Thermo Fisher Scientific, Waltham, MA) at a 1:10 dilution. Blot intensities were quantified using ImageJ software (version 1.54d) (Schneider et al., 2012), and relative intensities of Iba1 were normalized to β-Actin and plotted using GraphPad Prism 10.5.0.

### VEGF-Induced Endothelial Cell Migration

Primary HUVEC (Human Umbilical Vein Endothelial Cells) cells (PromoCell, Heidelberg, Germany) were seeded at 3x10^4^ cells/well in 70 µL endothelial cell media (PromoCell, Heidelberg, Germany) into a 2-well silicone insert (Ibidi, Munich, Germany) and incubated overnight at 37 °C with 5% CO_2_. Once the cells reached confluency, the silicone insert was removed to create a defined cell-free gap. The HUVEC cells were then stimulated with VEGF, a pro-angiogenic factor, along with BEM and MINO (100 µM) in the culture medium, which also contained other growth factors. The cells were allowed to grow in the presence of the treatments for 16 hours, and migration into the cell-free gap was monitored by photographing the cells under a microscope. The closure of the cell-free gap was quantified using the wound healing assay plugin in ImageJ1 (version 1.54d) software (Schneider et al., 2012).

### L-Glutamine-Induced ROS-Oxidative Stress

The antioxidant activity of BEM was assessed using the ROS-Glo H_2_O_2_ bioluminescent assay kit (Promega, Madison, WI) in a cell culture model. HUVEC cells were plated at a density of 2x10^4^ cells/well in a Falcon 96-well white flat-bottom plate and cultured overnight at 37 °C with 5% CO_2_ to 70% confluency. The cell suspension was adjusted to achieve the appropriate cell density, ensuring 80 µL of suspension per well. BEM and MINO were tested at concentrations of 30, 60, 90, and 120 µM, added in 20 µL of media, along with H_2_O_2_ substrate at a final concentration of 25 µM, per kit instructions. The cells were co-incubated with 10 mM L-glutamine (Thermo Fisher Scientific, Waltham, MA) for 4 hours, a concentration previously determined to be optimal (Tikka et al., 2001; Kraus et al., 2005b; Ola, 2022). Following incubation, 100 µL of ROS-Glo detection solution was added, and the mixture was incubated for an additional 20 minutes. Relative luminescence was measured using a Tecan Infinite M1000 PRO Monochromator Microplate Reader. The resulting luminescence was plotted as percentage control. The resulting concentration vs % control graphs were compared using simple linear regression analysis in GraphPad Prism 10.5.0.

### Anti-Inflammatory Activity

The anti-inflammatory activity of BEM was evaluated using the lipoxygenase inhibitor screening assay kit (Abnova, Taipei, Taiwan), following the manufacturer’s protocol. In brief, 15-LOX standard was treated with Nordihydroguaiaretic Acid (NDGA) as the positive control. BEM and MINO were tested as inhibitors at various concentrations (30, 60, 90, 120, and 200 µM) to assess the dose-response effect. The reaction was initiated by adding arachidonic acid substrate, followed by incubation on a shaker for 5 minutes. A chromogen mixture, provided by the manufacturer, was added to develop the reaction, which was incubated for an additional 5 minutes. Absorbance was measured at 490-500 nm using a plate reader. The average absorbance of all samples was determined, with the absorbance of blank wells subtracted from the 100% initial activity and inhibitor wells.

Percent inhibition was calculated as follows:

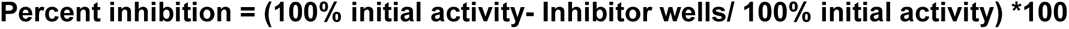

Dose-dependent inhibition curves were generated using non-linear regression analysis using a log(inhibitor) vs. response – variable slope (four parameters) model and comparisons between BEM and MINO were made using simple linear regression analysis in GraphPad Prism 10.5.0.

### Mitochondrial Toxicity

Mitochondrial toxicity of BEM was assessed using Human Colorectal Tumor (HCT) 116 cells (Mathur et al., 2024), as we determined this cell line to be sensitive to 10 μM of sodium azide. In a 96-well plate format, 200,000 cells/mL were plated in fresh glucose-containing DMEM media and galactose-containing DMEM medium. The cells were cultured overnight at 37 °C with 5% CO_2_. A working stock of 10 µM sodium azide (Sigma-Aldrich, S8032) was used as a control mitochondrial toxin to induce mitochondrial dysfunction in glucose and galactose-supplemented medium without serum. The cell culture conditions (such as using glucose and galactose) were optimized to enhance mitochondrial sensitivity. BEM and MINO were prepared for treatment at various concentrations (30, 60, 90, 120, and 200 µM) in both glucose and galactose-containing DMEM media.

The Mitochondrial ToxGlo™ Assay kit (Promega Corporation, Cat No. G8000) was used to measure cellular Adenosine Triphosphate (ATP) levels, comparing treated cells with vehicle controls after brief exposures to BEM and MINO. ATP levels were determined by adding an ATP detection reagent, which generates a luminescent signal proportional to ATP. The plate was allowed to equilibrate to room temperature (20 – 25 ^0^C) (USP, 2017) for 5 – 10 minutes before 100 µL of ATP detection reagent was added. After shaking for 1 – 5 minutes, luminescence was measured, adjusting the luminometer gain based on the untreated control signal. The percentage response for each test well was calculated by dividing its luminescence value by the average luminescence of the untreated control. The percent response was plotted for each compound and control against the concentration of BEM and MINO.

### Statistics

Statistical analyses were performed in, and graphs were generated by GraphPad Prism v10.5.0 software (Boston, MA). Data is presented as mean (± SEM), unless noted otherwise. Two-way ANOVA with Sidak’s *post hoc* multiple comparisons test was used for analyzing the antibacterial assay, antifungal study, microglial activation assay, VEGF–induced endothelial cell migration and mitochondrial toxicity assay. The log transformed data were used to perform nonlinear regression analysis to calculate IC50 values for cell viability using a three-parameter log(inhibitor) vs. response model, and for MMP-9, MMP-8, and 15-LOX inhibition assays, using a four-parameter variable slope model. Simple linear regression analyses of log-transformed data were performed between BEM and MINO in MMP- 9/8, 15-LOX and L-Glutamine-induced ROS-oxidative stress assays, to draw comparisons using slopes and coefficient of determination (fit) in GraphPad Prism 10.5.0.

## Results

### 10-Butyl Ether Minocycline

BEM is an analog of MINO and its chemical structure is shown in Fig 1. The 10-butyl ether group was substituted onto the phenolic hydroxy group on C-10 position of the tetracyclic ring.

### Antimicrobial activity

Across all tested concentrations (1.56 - 50 µg/mL), BEM exhibited a significantly smaller ZOI compared to MINO. We found that BEM had significantly lost its antimicrobial action against *E. coli* (*F* (1, 60) = 243.6, p < 0.0001, n = 6 biological replicates, main effect of drug) *and S. typhi* (*F* (5, 60) = 116.8, p < 0.0001, n = 6 biological replicates, interaction between drug and concentration) compared to MINO (Fig 2a and 2b).

**Figure 2.**
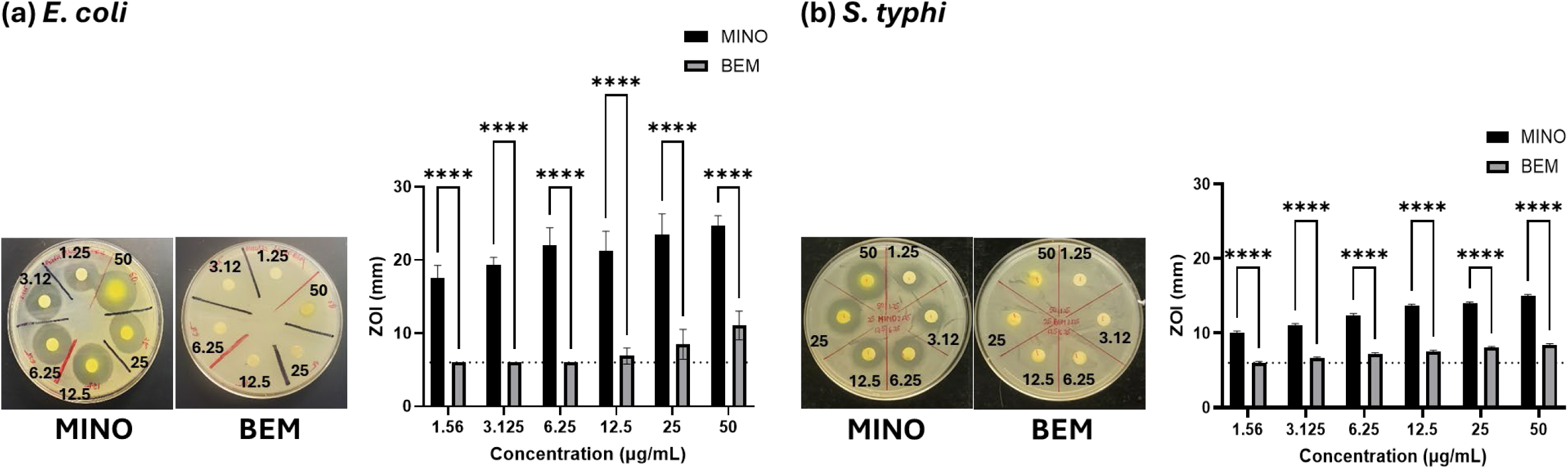
BEM lost antimicrobial action against *E. coli* and *S. typhi*. Zone of inhibition (ZOI) studies showed that BEM had a significant reduction in antibiotic activity against (a) *E. coli* and (b) *S. typhi.* Statistical analysis was performed using two-way ANOVA followed by Šídák’s *post hoc* multiple comparisons test. There was no interaction between concentration and drug in (a) while an interaction between drug and concentration was present in (b). Data are presented as mean ± SEM. Significance: ****p<0.0001; n = 6 biological replicates.

### Antifungal activity

BEM did not show a reduction in CFU of *C. albicans* (*F* (3, 16) = 248.2, p < 0.0001, n = 3 biological replicates), suggesting a loss of anti-fungal activity at all the tested concentrations (Fig 3). In contrast, MINO displayed significant anti-fungal activity at 50 and 100 μg/mL relative to BEM (Fig 3).

**Figure 3.**
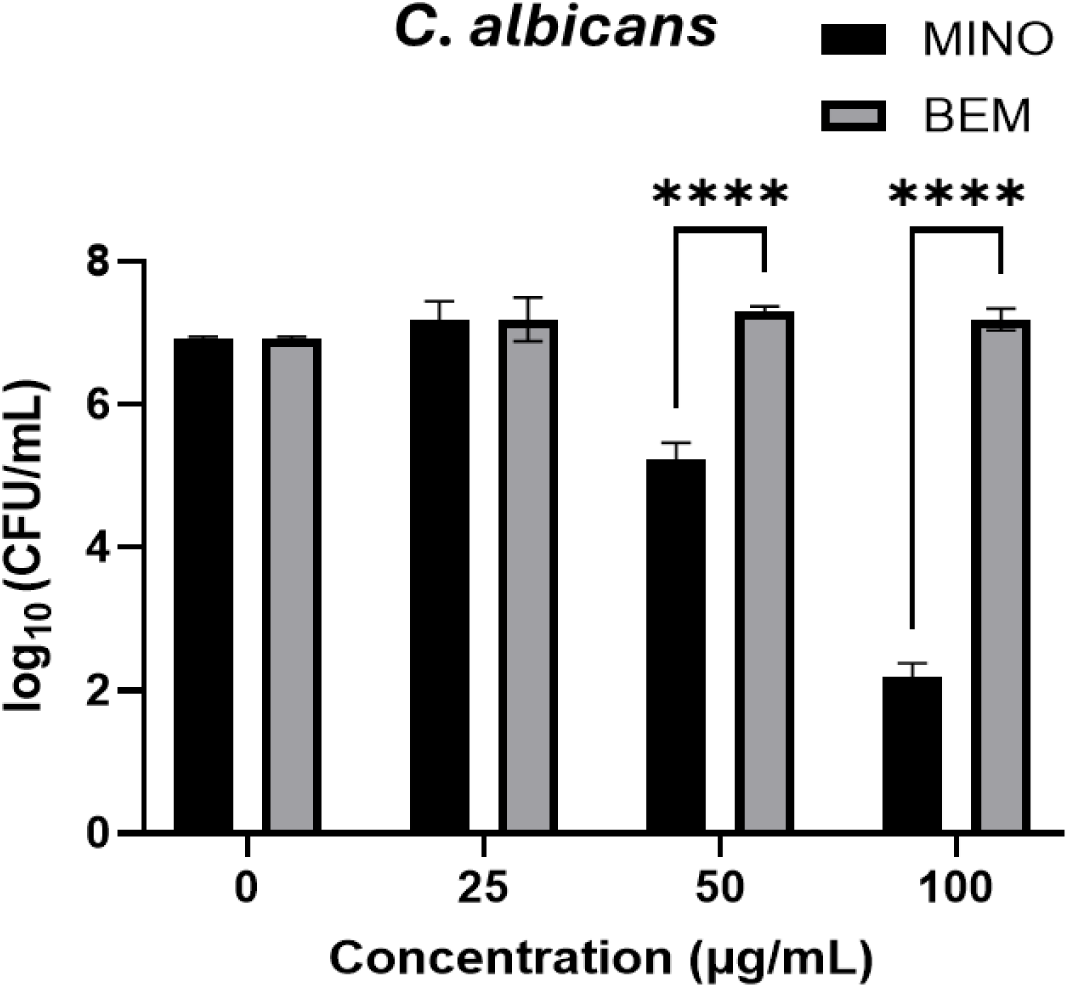
BEM did not show anti-fungal activity. BEM did not show a reduction in colony forming units (CFU) of *C. albicans* at all the tested concentrations, suggesting a loss of anti-fungal activity. MINO on the other hand showed significant antifungal action. Statistical analysis was performed using two-way ANOVA followed by Šídák’s *post hoc* multiple comparisons test. A significant interaction between concentration and drug was observed. Data are presented as mean ± SEM. Data are reported as mean ± SEM. Significance: ****p<0.0001 and n = 3 biological triplicates.

### BEM showed cell viability similar to MINO

The 24-hour cell viability of NB cells treated with BEM and MINO at the following concentrations: 0.1, 1.0, 125, 250 and 500 µM is shown in Fig 4a. Results are shown as percentage inhibition (n = 6 technical replicate wells per treatment). BEM, at low doses (0.1 – 10 µM), did not affect cell viability of NB cells, while BEM at high doses (≥50 µM), showed low cell viability. The IC50 of MINO was 43.2 µM and for BEM it was 102.1 µM, (Fig 4b and c) indicating that BEM is less cytotoxic, and tolerated at high concentrations than MINO.

**Figure 4.**
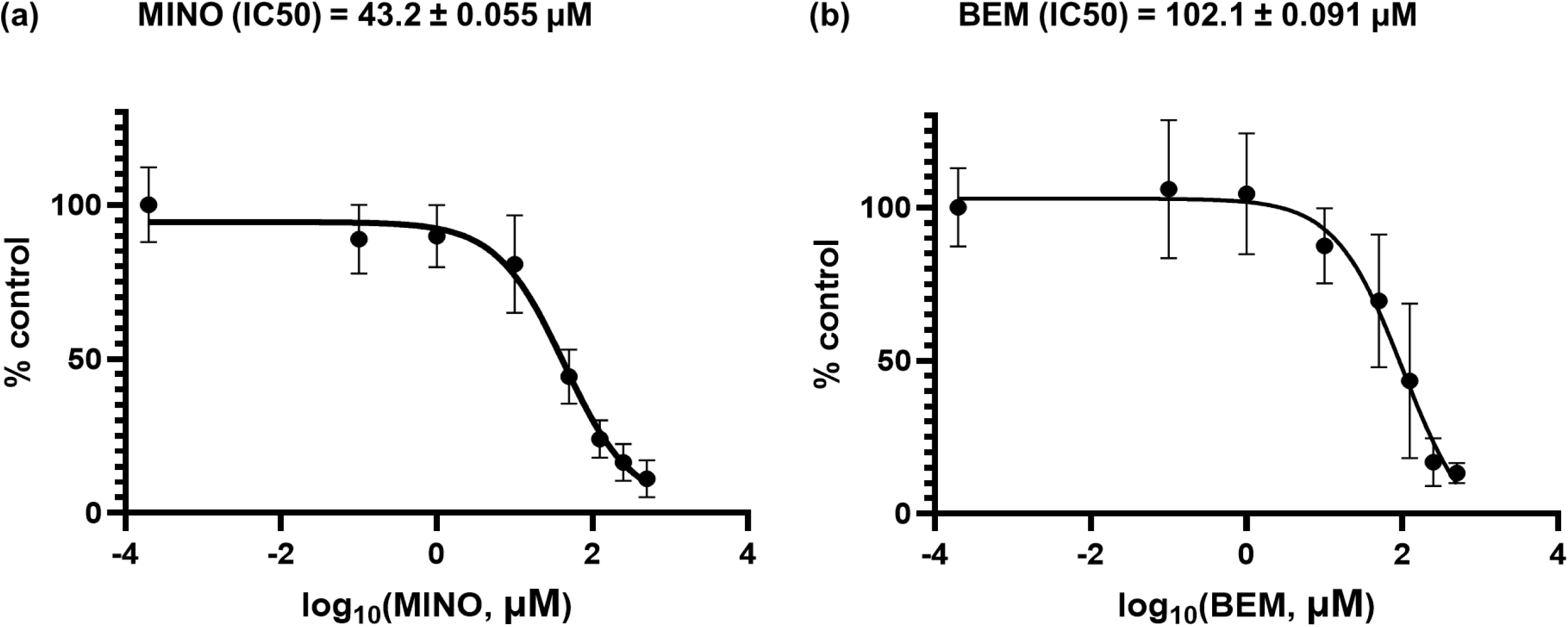
BEM demonstrated a dose-dependent reduction in NB cell viability, similarly to MINO. At low concentrations (0.1–10 µM), neither BEM nor MINO significantly affected NB cell viability. At higher doses (≥50 µM), both compounds showed cytotoxic effects. (a) The IC50 of MINO was determined to be 43.2 µM, and (b) the IC50 of BEM was 102.1 µM, calculated using a non-linear regression. These findings suggest that BEM could be tolerated at concentrations up to twice those of MINO. Data are presented as mean ± SEM from biological replicate wells; n = 6 per treatment.

### BEM and MINO inhibit MMP-9 and 8 in a concentration-dependent manner

The MMP-8/9 inhibitor screening assay was performed at serial concentrations of BEM and MINO, and compared against the reference inhibitor, NNGH. MMP-8/9 inhibitory action was calculated as percent inhibition. In Fig 5.1a, BEM’s inhibitory potency of MMP-9, was tested between 20, 40, 60 and 80 µM and BEM showed inhibition at 40, 60 and 80 µM. In contrast, MINO showed significant MMP-9 inhibition at 60 and 80 µM. Results are presented as percentage inhibition (n = 6 technical replicate wells). The IC50 of BEM for MMP-9 inhibition was 42.2 µM and for MINO, it was 60.3 µM (Fig 5.1b).

**Figure 5.1.**
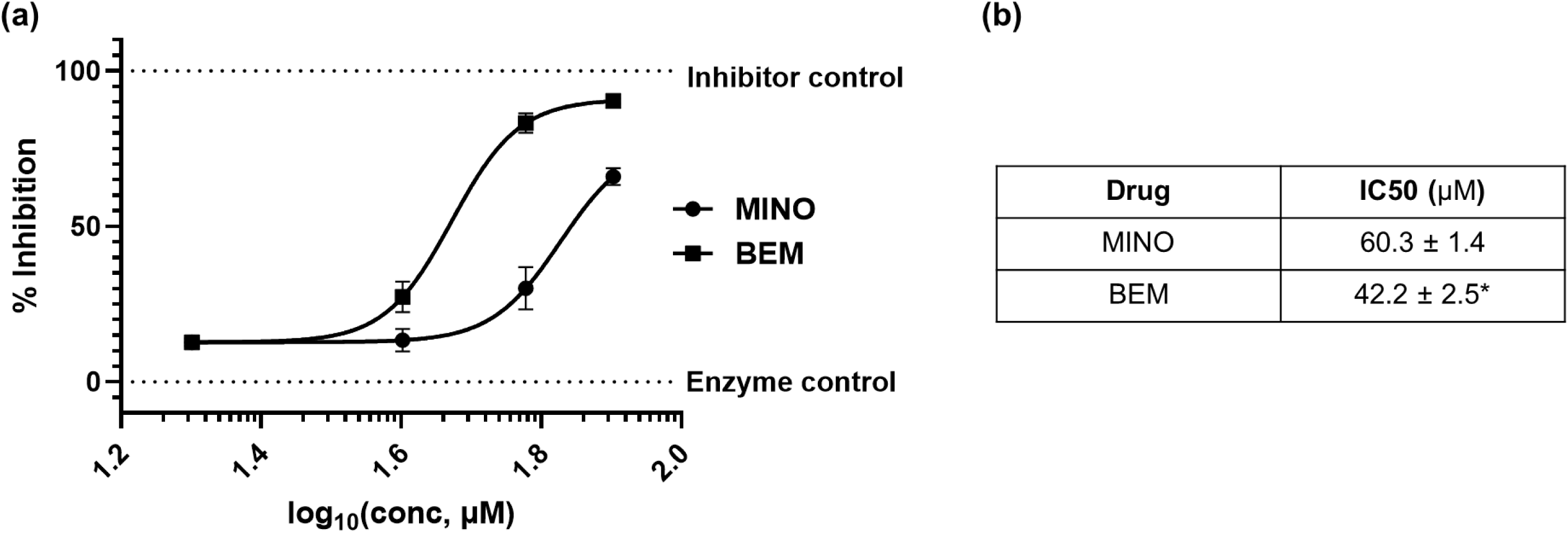
BEM exhibited higher potency than MINO in inhibiting MMP-9. The inhibitory potential of BEM and MINO against MMP-9 was evaluated in terms of percentage inhibition at various concentrations (20, 40, 60, 80 µM) in comparison to the prototypic inhibitor *N*-Isobutyl-*N*- (4-methoxyphenylsulfonyl)glycyl hydroxamic acid (NNGH). (a) Dose-dependent inhibition curves of MINO and BEM. The log transformed % inhibition v. concentration (µM) data was analyzed using non-linear regression to calculate IC50 values. (b) BEM exhibited a lower IC50 (47.0 µM; logIC50 = 1.67) compared to MINO (67.2 µM; logIC50 = 1.82), indicating higher potency. Regression analyses were restricted to the range of the data (i.e., from X = 1.477 to 2.079) and revealed a significant difference between the slopes of BEM and MINO (*p = 0.0180). The slope of BEM (m = 0.079 ± 0.13; r^2^ = 0.93) was significantly different than MINO (m = 0.14 ± 0.23; r^2^ = 0.72), suggesting that BEM and MINO differ significantly in their MMP-9 inhibitory potential. Data are shown as mean ± SEM from n = 5 technical replicate wells per concentration.

BEM’s inhibitory potency for MMP-8 was performed between 30, 60, 90 and 120 µM and the results (Fig 5.2a) are presented as percent inhibition (n = 5 technical replicate wells per treatment). The IC50 of BEM for MMP-8 inhibition was 69.4 µM and for MINO it was 45.4 µM (Fig 5.2b). Both MMP-8 and 9 showed a dose response inhibition to BEM and MINO. Higher concentrations (>200 µM) of BEM and MINO were not tested as the chromogenic nature of the tetracycline ring caused perturbations in absorbance measurements.

**Figure 5.2.**
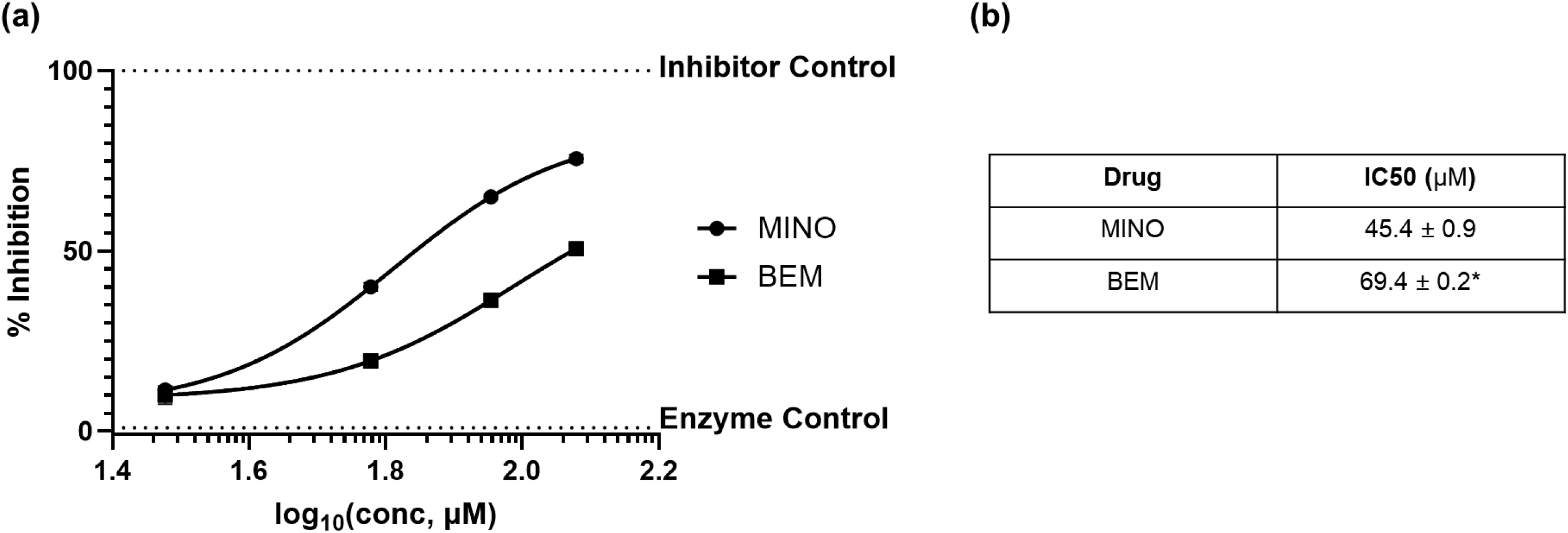
BEM and MINO showed MMP-8 inhibition in a concentration dependent manner. The inhibitory potential of BEM and MINO against MMP-8 was evaluated in terms of percentage inhibition at various concentrations (30, 60, 90, 120 µM) in comparison to the prototypic inhibitor *N*-Isobutyl-*N*-(4-methoxyphenylsulfonyl)glycyl hydroxamic acid (NNGH). (a) Dose-dependent inhibition curves of MINO and BEM. The log transformed % inhibition vs. concentration (µM) data was fit using nonlinear regression analysis to calculate IC50. (b) BEM’s IC50 (69.4 µM; logIC50 = 1.84) was higher compared to MINO (45.43 µM; logIC50 = 1.65). Regression analyses were restricted to the range of the data (i.e., from X = 1.477 to 2.079) and revealed significant difference between BEM and MINO (*p=0.0435). The slope of BEM (0.068 ± 0.12; r^2^ = 0.94) was significantly different than MINO (0.06 ± 0.12; r^2^ = 0.96), suggesting that BEM and MINO differ significantly in their MMP-8 inhibitory potential. Data are presented as mean ± SEM from n = 5 technical replicate wells per concentration.

### BEM suppressed LPS mediated microglial activation

Rat N9-microglial cells treated with LPS showed a change in phenotype relative to the control cells (Fig 6.1a). Iba1 IHC produced complete labeling to Iba1 localized in the cytoplasm of the rat N9 microglial cells. Results are presented as percentage ramified cells (*F* (3, 12) = 94.7, p < 0.0001, n = 6 biological replicates per treatment) (Fig 6.1b). The validity of IHC results were verified by performing western blot analysis of these N9-microglial cells (Fig 6.2a). The western blot analyses showed an upregulation of Iba1 protein in cells treated with LPS and a reduction of Iba1 in cells treated with BEM and MINO. Results are presented as normalized blot intensities (*F* (3, 8) = 31.8, p < 0.0001, n = 3 per treatment group, biological replicates from two independent experiments) (Fig 6.2b).

**Figure 6.1.**
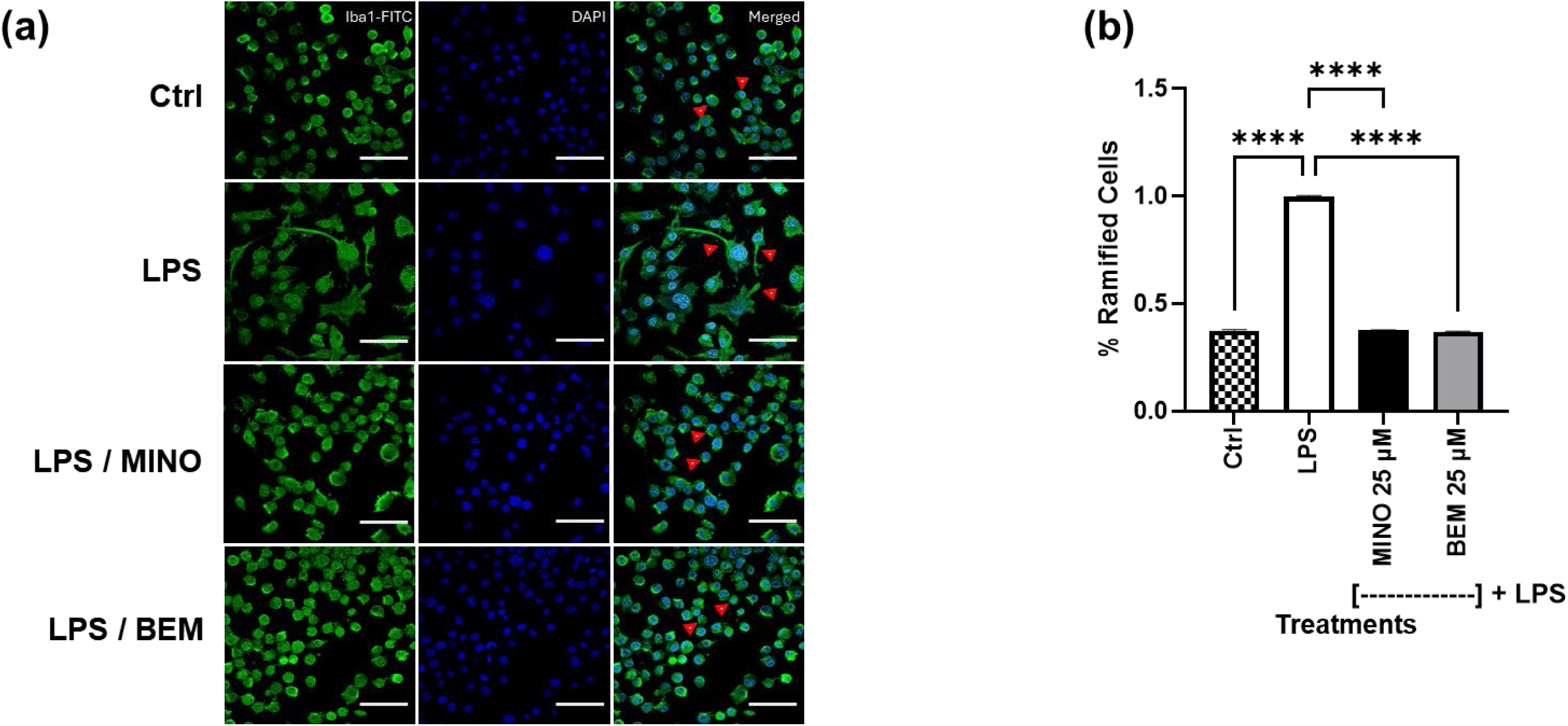
BEM suppressed LPS-mediated microglial activation in N9 microglial cells, similar to MINO. N9-microglial cells were treated with LPS (12 hours) followed by BEM or MINO (25 µM, 12 hours). (a) Immunocytochemistry of control and treatment groups. Nuclei were stained with DAPI, Iba1 was labelled with FITC; images were captured at 60X magnification. Red arrowheads: Ctrl: Amoeboid-like morphology of resting microglia. LPS: Ramified projections indicating reactive microgliosis. LPS + MINO / BEM: Amoeboid-like morphology was restored to a state comparable to the control group. (b) Quantification of % ramified cells. Data are expressed as mean ± SEM (n = 6 biological replicates per treatment). Statistical analysis was performed using two-way ANOVA followed by Šídák’s *post hoc* multiple comparisons test. Significance: ***p < 0.0001 (Ctrl vs. LPS, LPS vs. MINO 25 μM, LPS vs. BEM 25 μM); **p = 0.0014 (Ctrl vs. MINO 25 μM); **p = 0.0032 (MINO 25 μM vs. BEM 25 μM). Scale bar: 100 µm.

**Figure 6.2.**
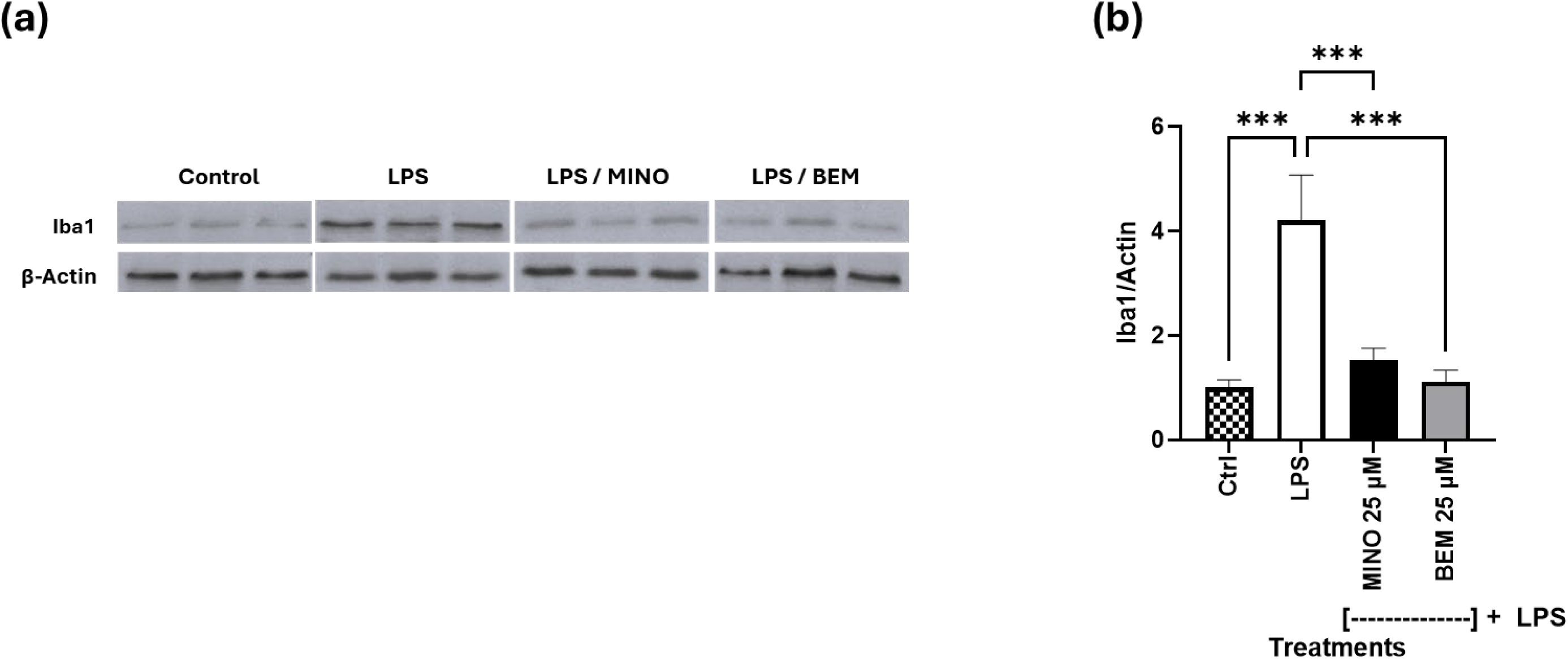
BEM reduced Iba1 expression in LPS-activated N9 microglial cells. N9 cells were treated with LPS for 12 hours, followed by 25 µM BEM or MINO treatment for an additional 12 hours. (a) Representative western blot showing basal Iba1 expression (control), upregulation after LPS, and reduction with BEM and MINO treatments. (b) Densitometric quantification of Iba1 levels normalized to actin. Data are presented as mean ± SEM (n = 3 per treatment group). Statistical analysis was performed using two-way ANOVA with Šídák’s *post hoc* multiple comparisons test. Significance: ***p = 0.0002 (LPS v Ctrl); ***p = 0.0006 (LPS V MINO 25 µM); ***p = 0.0002 (LPS V BEM 25 µM).

### BEM inhibited VEGF-induced endothelial cell migration

HUVEC cells were treated with VEGF and other growth factors, which induced endothelial cell migration. At lower concentrations (20, 40, 60 and 80 µM), BEM did not show anti-VEGF effect (data not presented). The effect on cell-migration at these lower concentrations was not significantly different from the positive control, VEGF. At 100 µM, BEM showed a measurable anti-VEGF effect (Fig 7a). Results are shown as area remaining (*F* (3, 12) = 15.3, p = 0.0002, n = 4 biological replicates per group) (Fig 7b).

**Figure 7.**
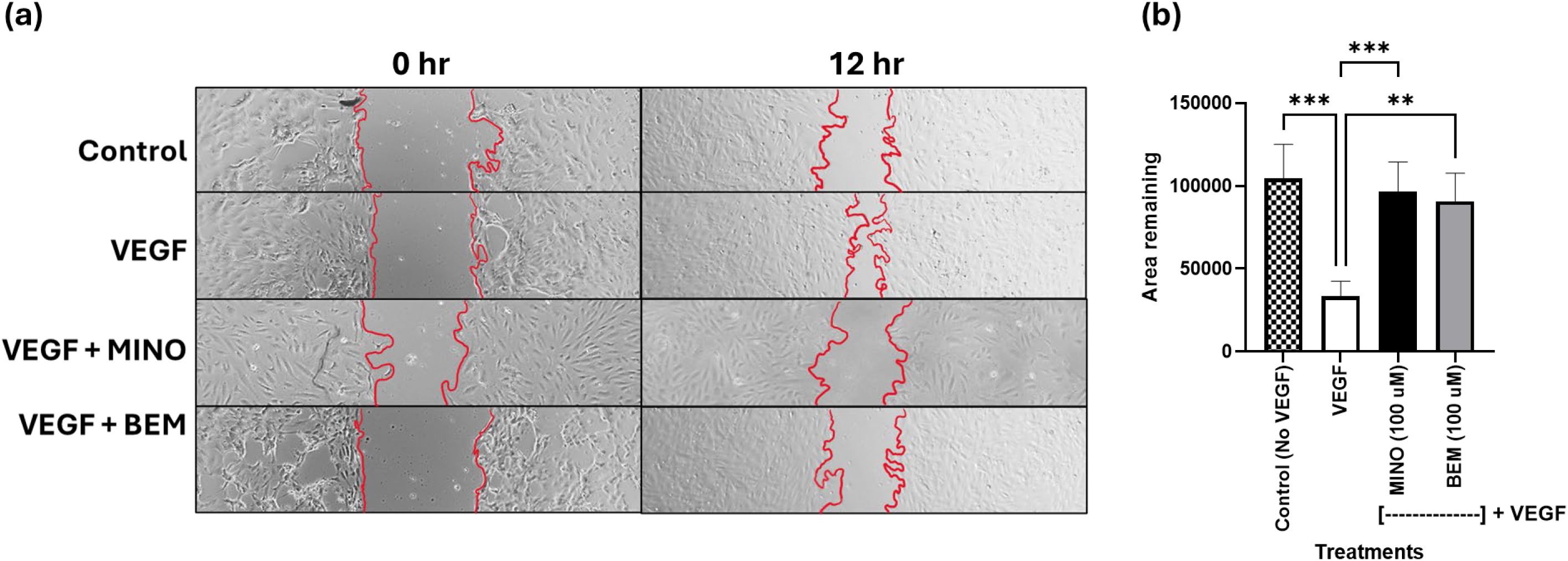
BEM inhibits VEGF-induced endothelial cell migration in HUVECs similarly to MINO. Human umbilical vein endothelial cells (HUVECs) were seeded at 3 × 10⁴ cells/well in endothelial cell medium into a two-well silicone insert to create a defined cell-free gap. After adherence, cells were treated with 25 ng/mL VEGF (positive control) or no VEGF (negative control), followed by treatment with either minocycline (MINO) or butyl ether minocycline (BEM) at 100 µM. (a) Representative phase-contrast images show VEGF-stimulated migration and its inhibition by BEM and MINO. (b) Quantification of the remaining gap area revealed significant inhibition of VEGF-induced migration by both BEM and MINO. Data are shown as mean ± SEM (n = 4 biological replicates per group). Statistical analysis was performed by two-way ANOVA with Šídák’s *post hoc* multiple comparisons test. Significance: ***p = 0.0003 (VEGF vs. Control), ***p = 0.0009, (VEGF vs. VEGF + MINO), **p = 0.0022 (VEGF vs. VEGF + BEM).

### BEM attenuated L-Glutamine induced ROS-oxidative stress

L-Glutamine induced ROS- oxidative stress was reduced by both BEM and MINO. We observed a dose-responsive reduction of ROS with both BEM and MINO. A significant reduction in ROS activity was observed at 60, 90 and 120 µM in both BEM and MINO (Fig. 8a). Results are shown as percentage control (n = 4 biological replicates per treatment).

**Figure 8.**
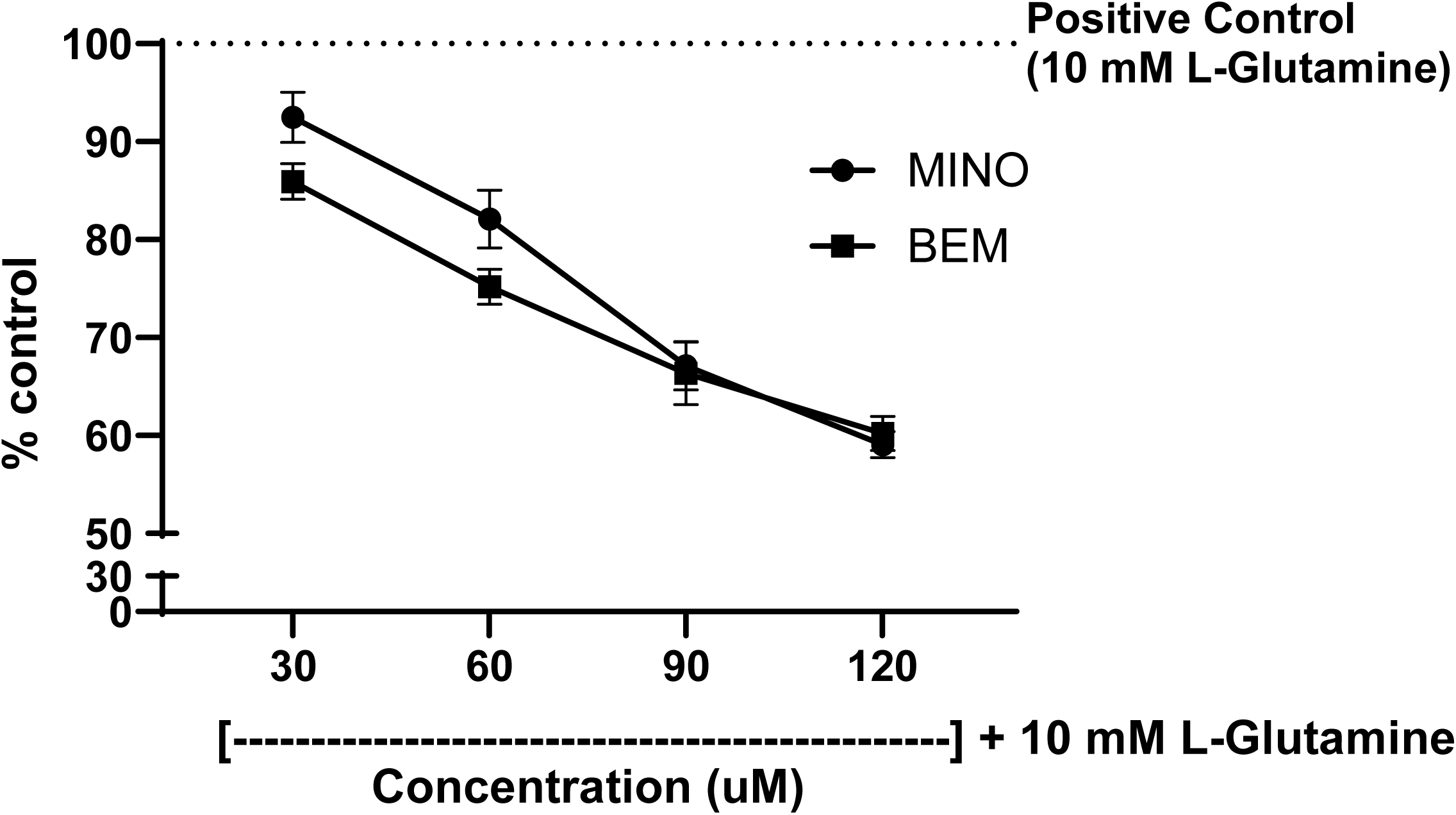
BEM and MINO attenuate L-glutamine–induced ROS production in a dose-dependent manner. N9 microglial cells were treated with 10 mM L-glutamine (L-Glu) for 12 hours to induce oxidative stress, followed by co-treatment with either butyl ether minocycline (BEM) or minocycline (MINO) at 30, 60, 90, or 120 µM. ROS levels were quantified and expressed as a percentage control. Using regression analyses, we can conclude that the differences between slopes of MINO (–0.32 ± 0.03; r² = 0.85) and BEM (–0.25 ± 0.03; r² = 0.84) is not significant (F = 2.181, p = 0.1509). However, both BEM and MINO effectively suppressed L-Glu–induced ROS within the tested dose range (60% of the control). Data are presented as mean ± SEM from n = 4 biological replicate wells per group.

### BEM and MINO showed moderate LOX inhibitory action

Both BEM and MINO showed a dose response effect compared to NDGA, a 5-, 12-, and 15-LOX inhibitor. Significant LOX inhibition was observed between 30, 60, 90, 120 and 200 µM for BEM and MINO (Fig. 9a). Results are plotted as percentage inhibition (n = 3 technical replicate wells per treatment). The IC50 of BEM for LOX inhibition was 92.6 µM and for MINO, it was 65.6 µM (Fig. 9b). At high concentrations (> 200µM), BEM and MINO had measurement interference due to chromogenic tetracyclic rings. The absorbances of treatments (60, 90, 120, 200 µM) were normalized to respective absorbances of pure drug compounds to account for this chromogenic effect.

**Figure 9.**
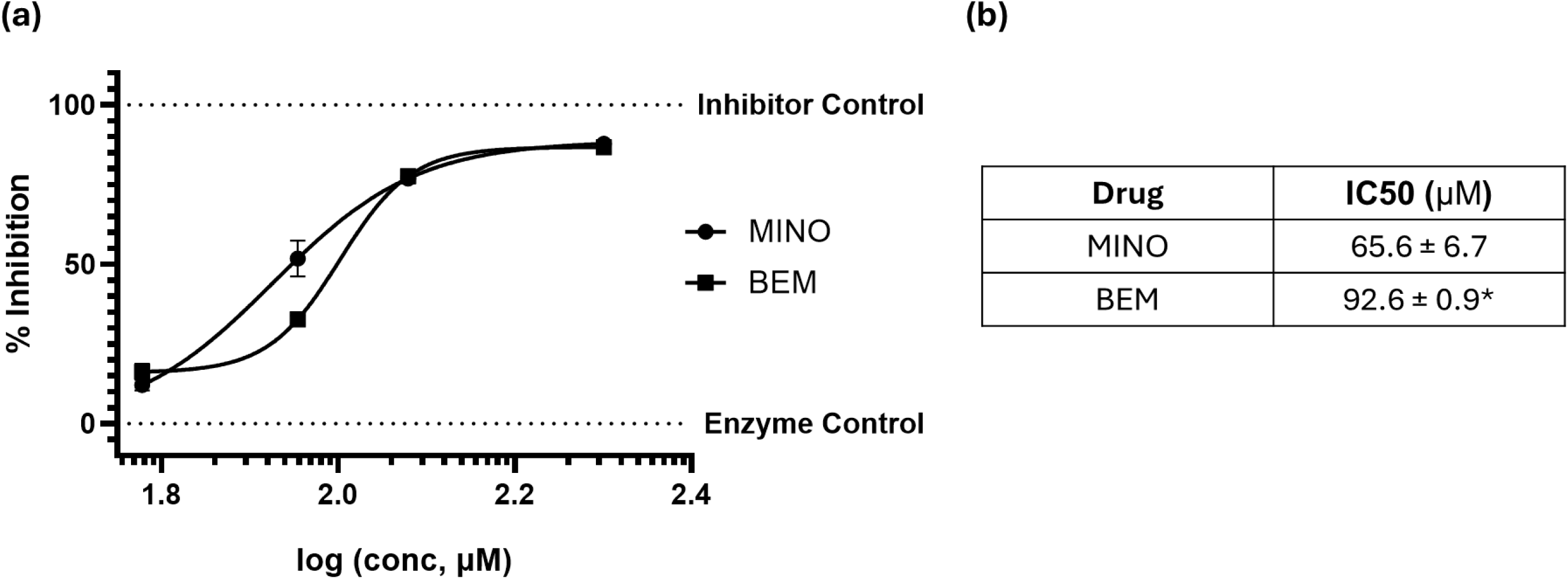
BEM and MINO showed modest LOX inhibitory action. The inhibitory potential of BEM and MINO against 15-LOX was evaluated across concentrations of 30, 60, 30 and 120 µM in comparison to the prototypical inhibitor, nordihydroguaiaretic acid (NDGA). (a) Dose-dependent inhibition curves of BEM and MINO. The log transformed % inhibition v. concentration (µM) data was fit using nonlinear regression analysis to calculate IC50. (b) The IC50 of BEM was 92.6 µM (logIC50 = 1.96) and for MINO it was 65.6 µM (logIC50 = 1.817). Regression analyses were restricted to the range of our data (i.e., from X = = 1.477; end at X = 2.079) and revealed no significant differences between the two treatments (*p = 0.6975). The slope of BEM (0.18 ± 0.37; r^2^ = 0.86) was not significantly different than MINO (0.27 ± 0.55; r^2^ = 0.76). Data are presented as mean ± SEM of n = 3 technical replicate wells per treatment.

### BEM is mitochondrial-safe

BEM and MINO did not alter the ATP levels in HCT-116 cells compared to the positive control NaN₃ (sodium azide) at various concentrations (30, 60, 90, 120, and 200 µM). ATP levels were measured under two conditions: cytoplasmic ATP (glucose) and mitochondrial ATP (galactose).

In both glucose and galactose conditions, NaN₃ significantly reduced ATP levels relative to all treatments. BEM and MINO maintained significantly higher ATP levels at all tested concentrations compared to NaN₃, indicating preserved mitochondrial function. BEM and MINO did not impact cytoplasmic or mitochondrial ATP production (Fig. 10). At the highest concentration (200 µM), ATP production was reduced by 68 % with BEM and 59 % with MINO (*F* (11, 96) = 218.4, p < 0.0001, n = 5 per treatment, biological triplicates, interaction between drug and concentration). The chromogenic properties of the tetracyclic rings in both BEM and MINO interfered with ATP measurement at concentrations above 200 µM.

**Figure 10.**
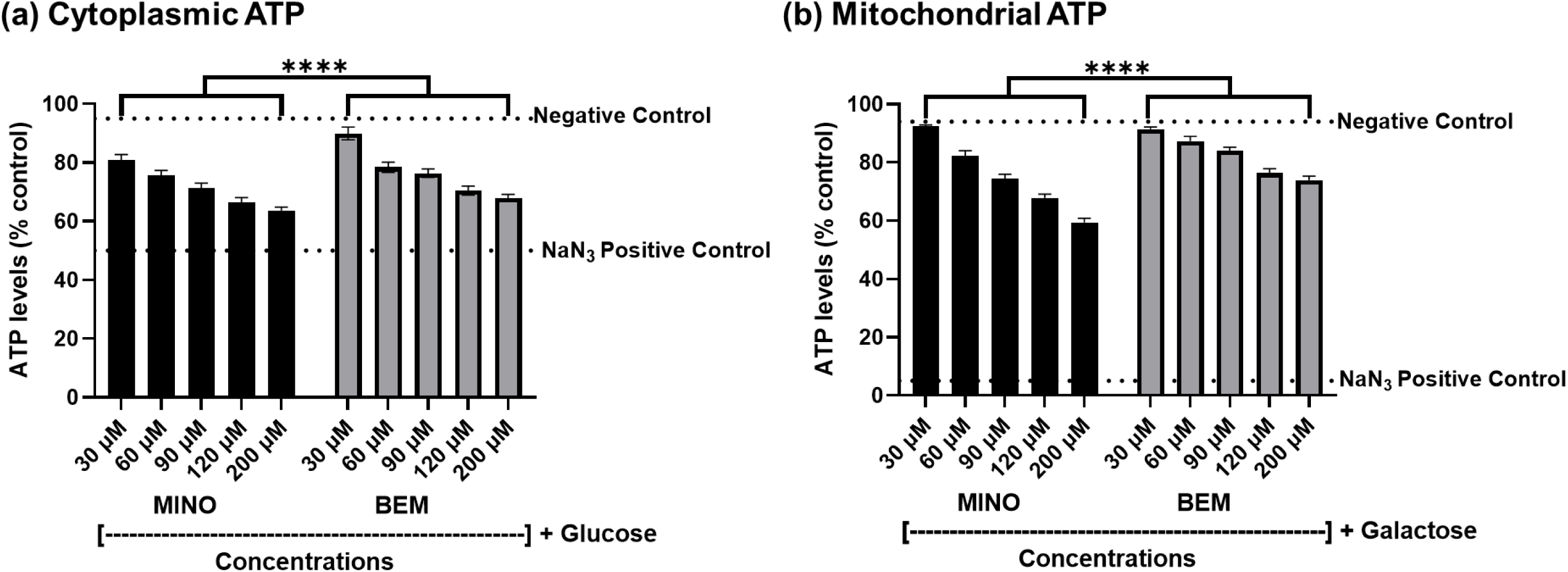
BEM and MINO do not exhibit significant mitochondrial toxicity compared to sodium azide. ATP levels were measured in Human Colorectal Tumor (HCT) 116 cells treated with MINO and BEM with increasing concentrations (30, 60, 90, 120, and 200 µM) in the presence of either glucose (Glu) or galactose (Gal) to distinguish cytoplasmic and mitochondrial ATP production, respectively. Sodium azide (NaN₃) served as the positive control for mitochondrial toxicity. Statistical analysis was performed using two-way ANOVA with Šídák’s *post hoc* multiple comparisons test. Data are presented as mean ± SEM of technical replicate wells (n = 5 per treatment, biological triplicates). ****p < 0.0001 for all comparisons between NaN₃ and other treatments (Ctrl, MINO, and BEM at all doses) under both Glu and Gal conditions.

## Discussion

The results of this study confirmed our hypothesis that BEM retained the known off-target actions and pharmacodynamic pleiotropy of MINO. The lack of anti-microbial activity of BEM against *E. coli, S. typhi* and antifungal action against *C. albicans* can be attributed to the modifications made at C-10 with a butyl substitution on MINO. MINO, on the other hand, showed a clear dose- dependent antimicrobial action. The C-10 position of tetracyclines, along with C-1, -11, and -12 are responsible for contact with the 30S rRNA in bacteria (Fuoco, 2012), which leads to suppression of protein synthesis and eventually, death. We proposed that the bulky C-10 butoxy group of BEM precludes a lock and key fit for the bacterial ribosome. However, it must be noted that other modifications could also result in changes in antimicrobial action. We chose to create BEM due to an *a priori* hypothesis that the butoxy moiety would a) preserve the pharmacophore responsible for the non-antimicrobial activities and b) increase lipophilicity, thereby, improving blood-brain barrier permeability for application in neuroimmune conditions. Of course, any structural modification to a drug can cause unexpected changes in mechanism(s) of action and side effects.

We observed a dose-dependent reduction in cell viability with BEM treatment. Interestingly, the calculated IC50 value indicated that BEM had improved cell viability, and was nearly half as cytotoxic as MINO. This result suggested that BEM may be well tolerated at higher concentrations than MINO. If such high tolerances translate to humans, BEM may be safer for long-term therapeutic use.

Our results confirmed that BEM preserved all MINO’s established, non-antimicrobial, mechanisms of actions that we tested, including the following: BEM was effective in inhibiting MMP-9 in a dose dependent manner. Since MMP-9 has been linked to alcohol reward and memory processes (Go et al., 2019; Valeri et al., 2023; Yin et al., 2023), and upregulation of MMPs is a marker of alcohol- induced damage in early stages of AUD, then inhibition of extracellular matrix proteins may be essential for restoration of synaptic plasticity related to addiction memory. For instance, MMP-9 inhibition blocked withdrawal-triggered escalation of alcohol self-administration in male Wistar rats, while neither altering locomotion nor producing persistent deficits in learning negative reinforcement (Go et al., 2019). In animal models, MMP expression and activity are closely linked to learning and memory, implicating MMPs in neuropsychiatric disorders such as addiction and depression (Beroun et al., 2019). MMP-8 is known to modulate inflammatory reactions in activated microglia and astrocytes (Lee et al., 2017). Though BEM demonstrated a lower MMP-8 inhibition compared to MINO, effective inhibition of MMP-8 by BEM at 120 µM was 50%, indicating a net inhibitory effect. Most tetracyclines, including MINO show MMP inhibition via a direct mechanism involving the chelation of divalent cations (like zinc, magnesium and calcium), which prevents MMPs from binding to other targets (Guerra et al., 2016). As such, we posit that BEM’s actions on MMPs occur through a similar mechanism, as we have not modified the functional groups responsible for MINO’s chelation of divalent cations.

Neuroinflammation has been implicated in various neurodegenerative and/or psychiatric disorders, including many substance use disorders. Microglia have been recognized as a significant participant in neuroinflammation. They are known to change between pro-inflammatory M1 and anti-inflammatory M2 states as a response function to various stimuli. Iba1, a calcium- dependent binding protein is commonly used as a biomarker for microglial activation, and may represent a therapeutic target, as its downregulation could help mitigate the neuroimmune dysregulation associated with AUD or other disorders (Barnett et al., 2020; Portis & Haass-Koffler, 2020; Warden et al., 2020; Carlson et al., 2025). In our experiment, LPS enhanced the expression of Iba1, a cytoskeletal protein, and triggered morphological changes in N9 microglia, suggesting polarization to a pro-inflammatory M1 state, which is similar to the acute ethanol-induced stress observed in AUD (Walter et al., 2017). The ramified projections in LPS treated microglial cells are distinct from the untreated cells, which are ameboid like (Jung et al., 2023). After treatment with BEM and MINO, LPS exposed cells showed a reversal of ramified to ameboid phenotype (similar to control). BEM effectively suppressed LPS-mediated microglial activation comparable to MINO- LPS treatment. The downregulation of Iba1 by BEM and MINO may play a crucial role in neuroimmune signaling and the allostatic sensitization of the stress circuitry (Koob et al., 2004; Koob, 2013). The Iba family of proteins are known to be localized in cellular projections and adhesion structures (Walter et al., 2017). We observed that LPS-treated N9 microglia exhibited prominent accumulation of Iba1 protein within cellular projections and adhesion structures, as shown by immunohistochemistry. Western blot quantification of Iba1 supported the immunohistochemistry findings, showing an effective reduction in Iba1 expression. We hypothesize that BEM chelates divalent cations, thereby contributing its direct inhibition of Iba1 analogous to inhibitory effects of MMPs. This hypothesis, however, requires a thorough investigation for BEM’s divalent cation requirement, which is beyond the scope of current study.

The inhibition of VEGF-induced migratory effects observed in HUVEC cells treated with BEM and MINO likely result from direct VEGF inhibition or by suppression of MMP action (Ramamurthy et al., 2002; Yao et al., 2004; Machado et al., 2006; Lee et al., 2007; Chang et al., 2017). By inhibiting cell migration, BEM may reduce inflammation and associated tissue damage, thereby contributing to neuroprotective effects in the central nervous system (J. M. Plane et al., 2010; Aghajani Shahrivar et al., 2022). Further investigation of specific pathways is needed to determine whether this mechanism is due to direct anti-VEGF action or direct inhibition of MMPs.

Glutamate-induced oxidative stress is a key factor in many neuroimmune and inflammatory disorders, including AUD (Roman & González, 2024). Glutamate, critical for cognition and mood, regulates memory, fear, and emotional processing in brain, specifically, in the hippocampus, prefrontal cortex, and amygdala (Nicoletti et al., 2023). In general, the process of addiction disrupts glutamate homeostasis, causing excess glutamate (excitotoxicity), elevated calcium levels, mitochondrial dysfunction, oxidative stress, and eventual neuronal atrophy and cell death (Nicosia et al., 2024). MINO had been demonstrated to act as a phenolic antioxidant (Kraus et al., 2005b). It had been proposed that the phenolic ring reacts with the free radical (ROS) and forms a phenol-derived free radical that is resonance stabilized. This specific stabilization effect was attributed to the C - 7 dimethyl amino substituent. In this study, BEM and MINO demonstrated near equivalent dose-dependent reductions in glutamine-induced ROS.

The LOX pathway plays a role in oxidative stress and inflammatory responses driven by microglia. (Chen & Zou, 2022). Under inflammatory conditions, 15-LOX is predominantly localized in the cytosol of the M1, proinflammatory, stage of microglia (Werz et al., 2018). The IC50 (92.6 µM) indicates that the anti-inflammatory action of BEM on LOX-15 is modest in potency. Further elucidation is needed to determine whether BEM can affect the downstream receptor action of LOX-15 enzymatic products, and whether its action includes inhibition of other family members such as LOX-5, and LOX-12. BEM’s inhibition of LPS-induced Iba1 expression, along with ROS inhibition and 15-LOX mediated anti-inflammatory effects in M1 microglia, likely contributes to its neuroprotective effects.

At high concentrations some early tetracyclines were known to impair eukaryotic mitochondrial function by affecting mitochondrial translation, which is evolutionarily conserved from endosymbiotic bacteria, leading to changes in cellular metabolism (Chatzispyrou et al., 2015; Moullan et al., 2015). Such subtle disruptions in mitochondrial function by tetracyclines can lead to disease tolerance mechanisms including tissue repair and metabolic reprogramming (Colaco et al., 2021; Sauer et al., 2022). BEM and MINO, tested at various concentrations against sodium azide-transmembrane potential depletion, did not show mitochondrial toxicity, even at concentrations higher than used in our previous animal studies (Liu et al., 2023; J. Willms et al., 2023; Liu, Gutierrez, et al., 2024; Liu, Panthagani, et al., 2024), reducing safety concerns from its long-term use.

The exact molecular pathways by which BEM shows pleiotropy are currently subject to investigation in our lab. Employing RNA sequencing data from alcohol and BEM treated C57BL/6J mice, the molecular mechanisms would be enumerated. We would like to report these findings in forthcoming communications. While few mechanisms have been established and well reported, this current study focuses on comparing BEM with those of established mechanisms of MINO.

Our data indicate that C-10 is not essential for maintaining the pleiotropic actions associated with MINO-like chemotype (BEM). Practically, C-10 can be treated as a modular position that can be diversified to block bacterial ribosomal binding yet leaving the C11/C12 β-diketone/phenolic region that underlies MMP engagement, redox buffering and microglial deactivation. We followed a Structure-Activity-Relationship modification axes to develop this ‘pleiotropy preservation paradigm’, with two major goals including: 1) steric shielding of C-10 to abolish bacterial 30S engagement. Modifications that were made or considered for this position are: n-alkyl or branched analogs to raise local bulk around C-10 without perturbing the chelating domain. These modifications were considered to explore electron donating elements on this position to subtly shift pKa/divalent cation affinity, potentially optimizing for MMP engagement while maintaining mitochondrial safety, and: 2) improve polarity/logD balance by C-10 modification to enhance CNS uptake and adjust solubility/permeability. Results from this study establish structure activity principles for C-10 substitution/modification to decouple antimicrobial activity from therapeutic pleiotropy, enabling the design of non-antibiotic tetracyclines for long-term neuroimmune indications such as AUD.

In conclusion, our findings indicate that BEM successfully retained several of the known pleiotropic therapeutic effects of MINO while eliminating its antimicrobial activity and reducing cytotoxicity. BEM effectively suppressed LPS-induced microglial activation, inhibited MMP-8 MMP-9, and 15-LOX activity, as well as reduced endothelial cell migration. BEM decreased oxidative stress while maintaining mitochondrial safety. These attributes position BEM as a promising therapeutic candidate for AUD (Liu et al., 2023; Liu, Gutierrez, et al., 2024; Liu, Panthagani, et al., 2024) and other neuroimmune or inflammatory (J. Willms et al., 2023) conditions. By shifting the focus from high-affinity, single target-based drug development to a multimodal-pleiotropic approach, this study highlights the potential of BEM as a safer, long-term treatment option for complex neurobiologically-based diseases. Furthermore, we identified that C-10 is not part of the pharmacophore responsible for BEM’s pleiotropic actions, adding to the existing literature of tetracycline’s structure activity relationship studies. The insights gained from BEM’s non-antibacterial mechanisms inform the design of next-generation therapeutics paving the way for innovative drug discovery strategies aimed at addressing multifaceted pathologies with significant unmet medical needs.

## Authorship Contributions: CRediT

Abdul Shaik: Conceptualization, methodology, data curation, formal analysis, writing - original draft, writing – review & editing.

Praneetha Panthagani: Conceptualization, methodology, data curation, formal analysis, writing - original draft, writing – review & editing.

Dr. Susan E. Bergeson: Conceptualization, formal analysis, writing - original draft, writing – review & editing.

Dr. Jeremy D. Bailoo: Formal analysis, writing - original draft, writing – review & editing. Dr. Stephanie Navarro-Turk: Methodology and data curation.

Dr. Xiaobo Liu: Methodology and data curation. Jeremy Garza: Methodology and data curation. Monica Aguilera: Methodology and data curation. Jordan Sanchez: Methodology and data curation. Kushal Gupta: Methodology and data curation.

Dr. Bruce Blough: Resources. Dr. Ted W. Reid: Resources. Dr. Abdul Hamood: Resources.

Elliott Pauli: Conceptualization, resources and project administration.

## Data Availability Statement

All data supporting the findings of this study are included within the manuscript. The raw datasets, have been deposited in Figshare and are publicly available at the following DOI: https://figshare.com/account/articles/29387459. Further details or specific requests for data may also be directed to.

## Abbreviations

BEM: 10-Butyl Ether Minocycline
MTT: 3-(4,5-Dimethylthiazol-2-yl)-2,5- Diphenyltetrazolium Bromide
DAPI: 4′,6-Diamidino-2-Phenylindole
5- LOX: 5-Lipoxygenase
ATP: Adenosine Triphosphate
AUD: Alcohol Use Disorder
CFU: Colony Forming Units
COX-2: Cyclooxygenase-2
DSM-V-TR, 2022: Diagnostic and Statistical Manual of Mental Disorders, Fifth Edition – Text Revision
DMSO: Dimethyl Sulfoxide
DMEM: Dulbecco’s Modified Eagle Medium
ECL: Enhance Chemiluminescence
*Escherichia coli* (*E. coli*) and *Salmonella typhi* (*S. typhi*)
FITC: Fluorescein Isothiocyanate
FDA: Food and Drug Administration
GMP: Good Manufacturing Practice
HRP: Horse Radish Peroxidase
(HCT 116) Cells: Human Colorectal Tumor
HUVEC: Human Umbilical Vein Endothelial Cells
IHC: Immunohistochemistry
IC50: Inhibitory Concentration 50
IL: Interleukin
Iba1: Ionized Calcium Binding Adaptor Molecule 1
LPS: Lipopolysaccharide
LB: Lipoxygenase (15-LOX) Lysogeny Broth
MMP-8 and -9: Matrix Metalloproteinases
MINO: Minocycline
NNGH: N-Isobutyl- N-(4-methoxyphenylsulfonyl)glycyl Hydroxamic Acid
NB: Neuroblast Cells
NDGA: Nordihydroguaiaretic Acid
PBS: Phosphate Buffered Saline
PVDF: Polyvinylidene Fluoride
ROS: Reactive Oxygen Species
RH: Relative Humidity
SEM: Standard Error of Mean
VEGF: Vascular Endothelial Growth Factor
YPD: Yeast Peptone Dextrose
ZOI: Zone of Inhibition

---

**Supplementary Figure 1.**
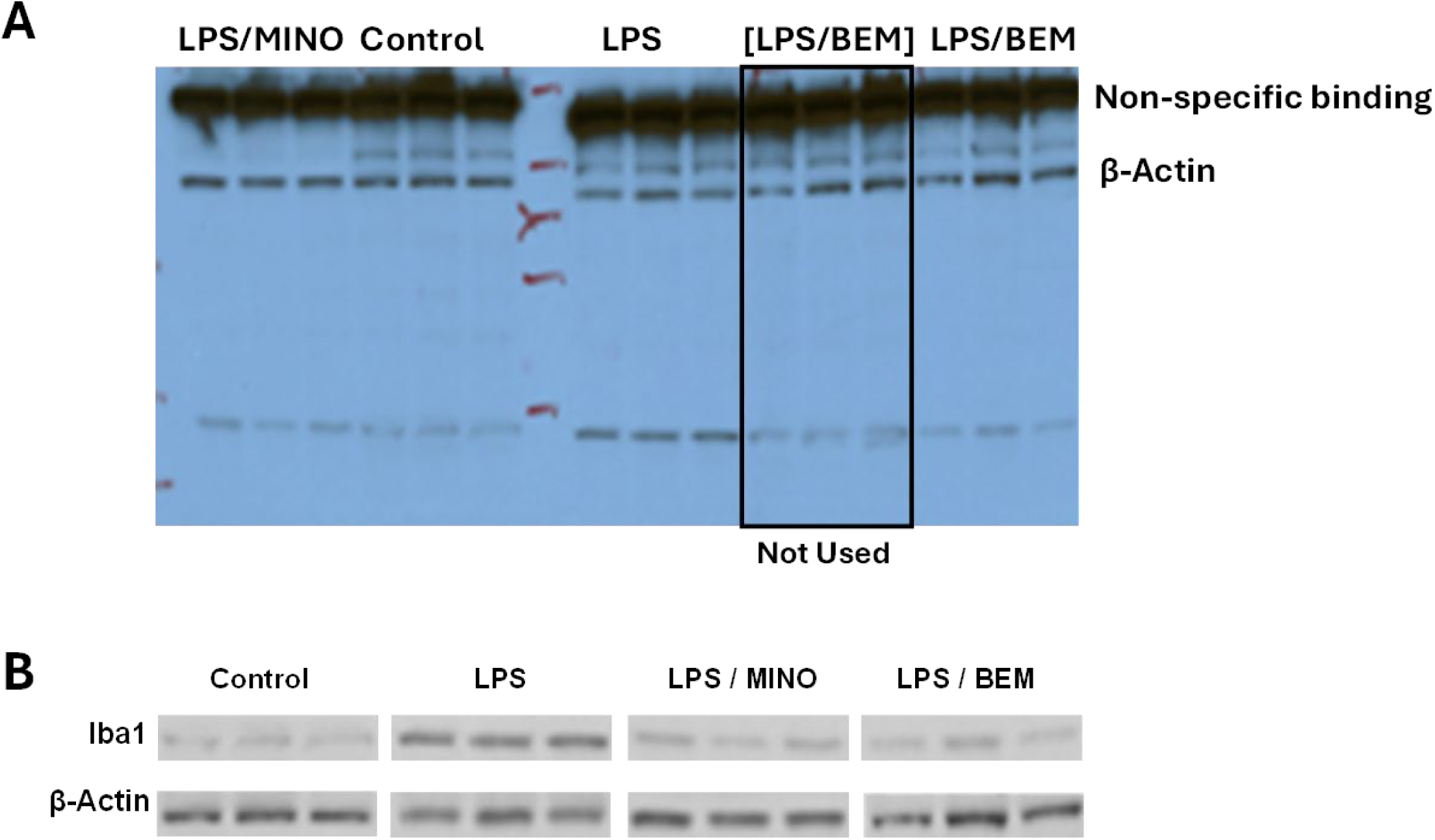
Original data for Figure 6.2. A) The western blot for the data from Fig 6.2. was exposed to Thermo Scientific CL-Xposure Film for 10 seconds. Note that three identical LPS/BEM samples were run twice. Those inside the black box were not used for the analysis. B) The film was converted to grayscale for densitometric measurements using ImageJ software (version 1.54d) and the display of individual treatment bands was performed using crop function in Microsoft Power Point by keeping the same cropping dimensions for all the proteins. The reporting order of treatments and proteins in B was reordered for logical consistency.

